# Long-term, cell type-specific effects of prenatal stress on dorsal striatum and relevant behaviors in mice

**DOI:** 10.1101/2024.12.27.627207

**Authors:** Maya M. Evans, Benjamin W. Q. Hing, Matthew A. Weber, Sara V. Maurer, Ahmed I. Baig, Grace S. Kim, Samantha L. Anema, Rhett M. Ellerbroek, Kartik Sivakumar, Jacob J. Michaelson, Nandakumar S. Narayanan, Hanna E. Stevens

## Abstract

Maternal stress during pregnancy, or prenatal stress, is a risk factor for neurodevelopmental disorders in offspring, including autism spectrum disorder (ASD). In ASD, dorsal striatum displays abnormalities correlating with symptom severity, but there is a gap in knowledge about dorsal striatal cellular and molecular mechanisms that may contribute. Using a mouse model, we investigated how prenatal stress impacted striatal-dependent behavior in adult offspring. We observed enhanced motor learning and earlier response times on an interval timing task, with accompanying changes in time-related medium spiny neuron (MSN) activity. We performed adult dorsal striatal single-cell RNA sequencing following prenatal stress which revealed differentially expressed genes (DEGs) in multiple cell types; downregulated DEGs were enriched for ribosome and translational pathways consistently in MSN subtypes, microglia, and somatostatin neurons. DEGs in MSN subtypes over-represented ASD risk genes and were enriched for synapse-related processes. These results provide insights into striatal alterations relevant to neurodevelopmental disorders.

## Introduction

Maternal stress during pregnancy, or prenatal stress, increases the risk for neurodevelopmental disorders (NDDs) in offspring^1-3^, including autism spectrum disorder (ASD) and attention-deficit hyperactivity disorder (ADHD). In multiple animal models, behavioral changes in offspring following prenatal stress have been associated with neurobiological changes, but mechanisms by which prenatal stress leads to these changes are not fully understood^4^. While risk associated with PS prenatal stress has smaller effect size on subsequent NDD diagnosis than some gene risks (e.g., meta-analysis odds ratio 1.64 for ASD, 1.72 for ADHD)^2^, the potential convergence of mechanisms across genetic and environmental factors may reveal important mechanisms for treatment targeting.

One commonality across NDDs in humans is abnormalities of dorsal striatum^5^. In children with ADHD, there is decreased functional connectivity between dorsal striatum and the cortex^6^ and reduced dorsal striatal size that does not persist to adulthood^7^. In ASD, dorsal striatum is enlarged and has increased functional cortical connectivity^8,9^. Dorsal striatum is also changed on cellular and molecular levels. Increased striatal dopamine receptor D2 (*Drd2*) transcript levels occur in post-mortem brain tissue from individuals with ASD^10^, and increased striatal Drd2-expressing cells are found in the 16p11.2 hemi-deletion ASD mouse model^11^. Disorganization of striatal striosome and matrix compartments occur in ASD^12^, as well as in the valproic acid mouse model of ASD^13^. Disrupted striatal glutamate dynamics also appear in both human ASD and ADHD, as well as in mouse models^11,14-17^. In addition, some defining behavioral features of ASD, including restricted, repetitive behavior, may be viewed as atypical expressions of striatal function^18^.

Changes in striatal-dependent learning and behavior have been observed in offspring after prenatal stress. One study investigated learning strategies in a navigation task in human subjects whose mothers had experienced a major negative life event during pregnancy^19^. The group exposed to prenatal stress more often used a rigid, response learning strategy, associated with the caudate, rather than a flexible, spatial learning strategy. Pregnant rats administered corticosterone (CORT), rodent stress hormone, during pregnancy had hyperactive juvenile offspring^20^. Additionally, repetitive restraint stress in pregnant mice impacts reversal learning, increases self-grooming, and impairs memory of a motor learning task^21,22^. These changes in learning and memory, activity level, and repetitive behavior are highly relevant to NDDs, suggesting that prenatal stress could have an etiological role. These studies also emphasize the susceptibility of the developing striatum, a substrate for these behaviors, to stressors during the prenatal period.

Prenatal stress in animal models also causes long-term changes in neurotransmitter dynamics in dorsal striatum. Positron emission tomography of rhesus monkeys exposed to prenatal stress revealed increased striatal dopamine transporter (DAT) availability, correlated with tactile hyperresponsivity^23^, a trait common in those with NDDs. For pregnant rats administered CORT, juvenile offspring show higher striatal dopamine metabolism^20^. Adult offspring of rats exposed to repetitive restraint stress during pregnancy also show increased expression of Drd2 in ventral striatum and increased NMDA expression in dorsal and ventral striatum^24^. Changes in striatal dopamine and glutamate dynamics following prenatal stress may contribute to changes in striatal-dependent learning and behavior in offspring. However, it remains unknown how prenatal stress affects specific striatal cell types.

Single cell RNA-sequencing (scRNAseq) is a powerful tool that can reveal transcriptional characteristics on a cell-type specific level. Understanding how specific cell types are affected or unaffected by risk factors like prenatal stress PS can help unveil mechanisms behind such risks. scRNAseq has been used to define the transcriptomic profiles of dorsal striatal cell types in wild type mice^25-28^ and in a Huntington’s disease mouse model^29^. As for mouse models of ASD, dorsal striatal single-nucleus RNA sequencing in the 16p11.2 deletion model has revealed changes in synaptic plasticity in dopamine receptor D1 (Drd1) and Drd2 MSNs and changes in GABAergic signaling in Drd1 MSNs^30^. Bulk RNA sequencing of dorsal striatum across single gene models of ASD has uncovered changes in several domains, including glutamatergic signaling and dopaminergic cell development^16,31-33^.

The effects of prenatal stress on the dorsal striatal transcriptome have not been explored in bulk striatal tissue, nor on the single cell level. Here, we employ dorsal striatal scRNAseq to investigate this gap in knowledge. With a mouse model, we used a repetitive restraint stress paradigm from embryonic day 12 through birth, a critical window of neurogenesis, gliogenesis, and cell fate determination in striatal development^34^. When offspring reached adulthood, we performed behavioral tests relevant to dorsal striatal functions. We also performed single unit recordings from dorsal striatum during behavior to capture functional changes at multiple levels. In separate littermates, we collected dorsal striatum for scRNAseq. We found significant changes in striatal-dependent behavior following prenatal stress with accompanying changes in striatal neuron activity. Alongside this, we saw similar changes in the striatal transcriptome of multiple cell types, especially in the domains of translation, ribosome biogenesis, and synapse function.

## Results

### Prenatal stress affected dorsal striatal-dependent behaviors

A 30-minute open field test was conducted to probe locomotor activity and anxiety-relevant behavior. Prenatal stress (PS) had no effect on distance traveled (t(40) = 0.986; p = 0.330, Fig. 1a) or time spent in the center zone (t(40) = 0.346; p = 0.731, Fig. 1b).

**Fig. 1:**
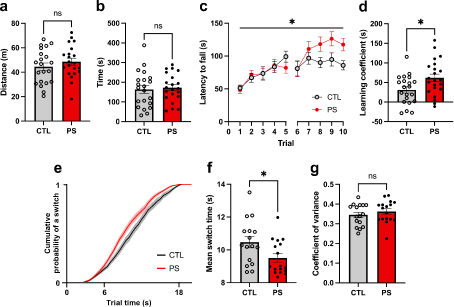
Prenatal stress (PS) resulted in persistent changes to motor learning and interval timing behavior. PS had no effect on (**a**) total meters (m) traveled or (**b**) seconds (s) in center during a 30-minute open field test. (**c**) PS increased latency to fall on accelerating rotarod across ten trials performed over two consecutive days beyond that shown for controls (CTL). (**d**) PS increased the learning coefficient (LC) from rotarod training. LC is calculated using the difference between the average of the last two trials and the average of the first two trials for each mouse. (**e**) Cumulative distribution function displaying probability of a switch as a function of time during long trials in the interval timing task averaged across CTL and PS mice. Shaded area is SEM. PS decreased (**f**) mean switch time, but did not alter (**g**) switch time coefficient of variance compared to CTL on the interval timing switch task. ns = non-significant. * = p<0.05. Error bars represent standard error of the mean.

To assess motor learning, mice were trained on an accelerating rotarod with five trials per day over two consecutive days. Latency to fall or second full rotation around the rod was recorded. There was a significant interaction between time and condition (F(9, 369) = 2.754; p = 0.004; Fig. 1c). PS mice displayed greater increase in latency to fall across time compared to control (CTL) (t(41) = 2.442; p = 0.019, Fig. 1d).

Body weight showed a trending negative correlation with average latency to fall on rotarod in males (r = -0.378; p = 0.082) and a significant negative correlation in females (r = -0.622; p = 0.003). However, body weight did not differ between the PS and CTL groups at adulthood in either males (t(20) = 0.123; p = 0.903) or females (t(19) = 0.304; p = 0.764).

To assess interval timing behavior, we focused on mean switch time and switch time coefficient of variance (CV) from the interval timing switch task. PS mice displayed earlier mean switch times (t(31) = 2.268; p = 0.029, Fig. 1e,f). However, PS mice did not display differences in switch time CV (t(31) = 0.901; p = 0.374, Fig. 1g), number of switch trials completed, number of rewards received, or number of nosepoke responses (Supplementary Fig. 1a-c).

### Dorsal striatal time-related neuronal activity was altered after prenatal stress

Multielectrode array recordings from dorsomedial striatum were conducted during the interval timing switch task in CTL and PS mice. Activity was analyzed for all units that exhibited firing rates above 0.5 Hz. We focused our analyses on MSNs, which have an essential role in timing behavior^35-37^. Recordings yielded a total of 84 MSNs from CTL mice and 78 MSNs from PS mice. The activity of each neuron was analyzed during each long trial in which the mouse received a reward, i.e., the mouse successfully switched during the long trial.

Some MSNs exhibited time-related, monotonic changes in firing rate, or “ramping” activity, over the first six seconds of the trial during timing behavior^35-37^. The mean estimated firing rate across all trials for two example neurons that display ramping activity are shown in traces (Fig. 2a). In a principal component analysis of MSN activity, this ramping pattern was represented in the first principal component (PC1) (Fig. 2b), which explained 55.5% of the variance in patterns of MSN activity over the first six seconds of switch trials (Fig. 2c). The ramping pattern of activity in MSNs is the strongest neural predictor of how much time has objectively passed in a trial^37^, meaning that MSN activity during the first six seconds of a trial is most predictive of objective time^37^. Therefore, we focused our analyses on that period (Fig. 2d,e). At the ensemble level, the percentage of dorsal striatal ramping MSNs was significantly higher in the PS group (51%) than in the CTL group (19%) (p < 0.0001; Fig. 2f). Interestingly, we found increased absolute value PC1 scores in PS MSNs compared to CTL MSNs (t(160) = 2.693; p = 0.0078; Fig. 2g), suggesting that PS MSNs ramp faster over the first six seconds of switch trials. This is further supported by increased absolute value firing rate slope derived from trial-by-trial GLMs over the same period in PS MSNs compared to CTL MSNs (t(160) = 3.439; p = 0.0007; Fig. 2h).

**Fig. 2:**
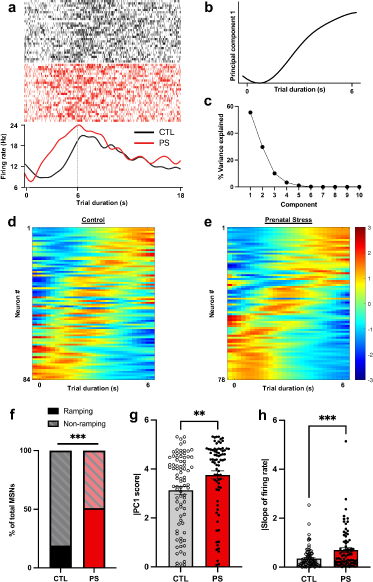
Prenatal stress (PS) altered striatal medium spiny neuron (MSN) activity during interval timing behavior. (**a**) Exemplar MSNs from one control (CTL) mouse and one PS mouse that show time-related ramping activity over the first six seconds of the trial. Each row in the raster plot represents one trial, and each tick mark represents an action potential. (**b**) The first principal component (PC1) represented time-related ramping activity over the first six seconds of the trial. (**c**) PC1 explained 55.5% of the variance in neuronal activity patterns. (**d-e**) Peri-event time histograms showing the Z-scored firing rate for each neuron in the CTL (**d**) and PS (**e**) ensembles. Each row represents an individual neuron. PS increased (**f**) the percentage of ramping MSNs at the ensemble level, (**g**) PC1 score (absolute value) in MSNs, and (**h**) the slope of firing rate (absolute value) in MSNs from 0–6 seconds. ** = p < 0.01. *** = p < 0.001. Error bars represent standard error of the mean.

### Multiple dorsal striatal cell types showed persistent changes in transcriptome after prenatal stress

scRNAseq analysis of dorsal striatum in 8–12-week-old mice showed clustered cell types by optimizing a modularity function and plotted using t-distributed stochastic neighbor embedding (tSNE) (Fig. 3a). From these clusters, striatal cell types were identified using gene expression markers (Supplementary Fig. 2,3,4), and clusters were not affected by batch or condition (Supplementary Fig. 5). Based on the clustering results, cell type proportions did not vary significantly between CTL and PS dorsal striatum after correction for multiple comparisons (Fig. 3b). PS induced a trend lower percentage of putative neuroblasts compared to controls (p = 0.050). Immunofluorescent stereology of intact sections from separate littermates aligned with this finding, showing no differences in Drd1 MSN, Drd2 MSN, overall GABAergic neuron, microglia, or astrocyte density in dorsal striatum (Supplementary Fig. 1f-k).

**Fig. 3:**
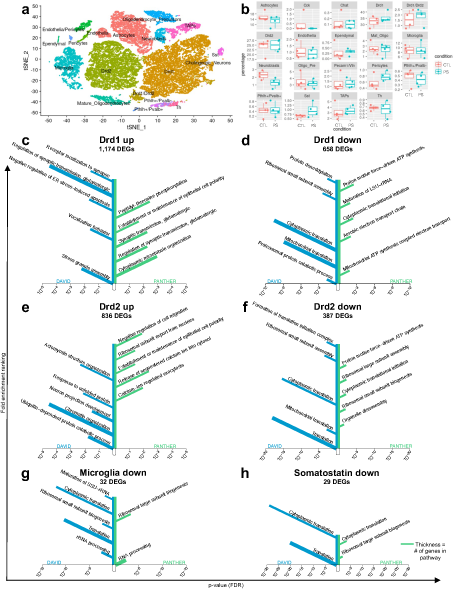
Prenatal stress (PS) altered transcription of genes related to synaptic function, ribosomes, and translation in multiple striatal cell types. **(a)** T-distributed stochastic neighbor embedding (tSNE) visualization of scRNAseq clusters of dorsal striatal cells (n = 31,040) from 8 samples (n = 4 control (CTL), 4 PS). (**b**) Box plots showing percentage of each cell type as a fraction of total cells in each sample with no differences between PS and CTL. (**c-h**) GO Tree Vis plots showing the top significantly enriched pathways among differentially expressed genes. x-axis represents false discovery rate (FDR); y-axis represents fold enrichment ranking; thickness of branch represents number of genes in reference pathway. Left side of chart shows results from DAVID; right side of chart shows results from PANTHER. Some GO terms were abbreviated: Negative regulation of ER stress-induced apoptosis ➔ Negative regulation of endoplasmic reticulum stress-induced intrinsic apoptotic signaling pathway Establishment or maintenance of epithelial cell polarity ➔ Establishment or maintenance of epithelial cell apical/basal polarity Maturation of LSU-rRNA ➔ Maturation of LSU-rRNA from tricistronic rRNA transcript (SSU-rRNA, 5.8S rRNA, LSU-rRNA) Formation of translation initiation complex ➔ Formation of cytoplasmic translation initiation complex

There were numerous differentially expressed genes (DEGs) by PS across striatal cell types (Supplementary Table 1, Fig. 3). The majority of DEGs were upregulated in PS. The most numerous subtypes, Drd1 and Drd2 MSNs, had substantial overlap of DEGs, with 503 shared DEGs being upregulated in both cell types (43% and 60% of each cell type DEGs, respectively) and 216 shared DEGs being downregulated in both cell types (33% and 56%, respectively) (Supplementary Fig. 2a,b). There were some shared DEGs between other cell types, as well (Supplementary Fig. 2a,b).

Drd1 and Drd2 MSNs were further subclustered, resulting in the identification of patch, matrix, and eccentric subtypes (Supplementary Fig. 2e,f). As expected, there were also numerous DEGs within these subpopulations (Supplementary Table 1).

Pathway enrichment analysis resulted in numerous significantly enriched pathways among DEGs for Drd1 MSNs, Drd2 MSNs, microglia, and somatostatin (Sst) interneurons, using both DAVID and PANTHER (Supplementary Table 1). Enriched pathways among upregulated DEGs in Drd1 and Drd2 MSNs showed common themes of synapse structure and function (e.g., “synaptic transmission, glutamatergic” and “neuron projection development”) and calcium signaling (e.g., “calcium-ion regulated exocytosis” and “release of sequestered calcium ion into cytosol”) (Fig. 3c,e). Enriched pathways among downregulated DEGs in both MSN subtypes as well as microglia and somatostatin (Sst) interneurons showed themes of translation (e.g., “cytoplasmic translation”; “translation”; and “cytoplasmic translational initiation”) and ribosome structure (e.g., “ribosomal large subunit assembly”; “ribosomal small subunit assembly”; “maturation of large subunit-rRNA”; and “maturation of small subunit-rRNA”) (Fig. 3d,f,g,h). Pathway enrichment analysis on DEGs from Drd1 and Drd2 matrix subpopulations also yielded significant results, which were highly similar to the results of the total Drd1 and Drd2 MSN populations. Other cellular subtypes, typically with small numbers of DEGs, demonstrated no significantly enriched pathways.

A comparison analysis using Ingenuity Pathway Analysis showed that canonical pathway results for Drd1 MSNs, Drd1 matrix MSNs, Drd2 MSNs, Drd2 matrix MSNs, microglia, and Sst interneurons were largely similar in content and directionality (Supplementary Fig. 2c) and included additional shared pathways for amino acid deficiency/metabolism, nonsense mediated decay, and the Rho GTPase cycle across cell types.

### Differential gene expression in striatum after prenatal stress overlapped with ASD-but not ADHD-associated genes

ASD-associated genes from the SFARI Gene database were significantly overrepresented in DEGs of both Drd1 (p = 8.59E-23) and Drd2 (p = 3.30E-11) MSNs after PS (Fig. 4a,b). Pathway analysis on ASD-associated genes among the Drd1 DEGs showed terms including “synapse assembly” and “histone H3-K9 demethylation” among others (Fig. 4c). The same analysis using ASD-associated genes among the Drd2 DEGs resulted in similar terms (Fig. 4d). A curated list of candidate ADHD risk genes was not overrepresented in DEGs of Drd1 (p > 0.999) or Drd2 (p = 0.337) MSNs.

**Fig. 4:**
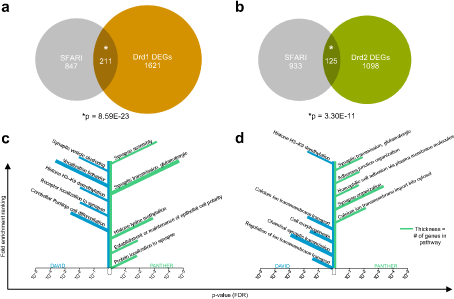
Prenatal stress (PS) altered ASD-associated genes in dorsal striatum. Venn diagram showing overlap between genes from the SFARI Gene database and differentially expressed genes in (**a**) Drd1 and (**b**) Drd2 MSNs after PS. (**c-d**) GO Tree Vis plots showing significantly enriched pathways among SFARI genes that were differentially expressed after PS. x-axis represents false discovery rate (FDR); y-axis represents fold enrichment ranking; thickness of branch represents number of genes in reference pathway. Left side of chart shows results from DAVID; right side of chart shows results from PANTHER. Some GO terms were abbreviated: Establishment or maintenance of epithelial cell polarity ➔ Establishment or maintenance of epithelial cell apical/basal polarity Homophilic cell adhesion via plasma membrane molecules ➔ Homophilic cell adhesion via plasma membrane adhesion molecules

## Discussion

Prenatal stress confers increased risk for NDDs, including ASD, ADHD, and others. Dorsal striatum is altered in these disorders at various levels, molecular to morphological. Limited animal studies suggest dorsal striatum may be a substrate of PS-induced changes, but no studies have examined its transcriptional effects on dorsal striatum or specific dorsal striatal cell populations. For these reasons, we investigated effects of PS on dorsal striatum using a mouse model. We used repetitive restraint stress to model PS, administered on embryonic days 12–18 to target early striatal cell development. Then, we tested adult offspring on striatal-dependent behaviors, finding enhanced motor learning and earlier response times related to interval timing in PS offspring without gross changes to locomotor activity, confirming that this prenatal insult resulted in long-lasting changes relevant to striatum and neurodevelopmental-relevant outcomes. Next, we employed a multi-electrode array to record dorsal striatal neuronal activity in mice performing the interval timing task, observing a higher proportion of ramping MSNs and faster ramping in PS mice. Lastly, we performed scRNAseq on the dorsal striatum from adult male offspring and saw significant transcriptional changes, overlapping across many striatal cell types.

In previous work, maternal immune activation during pregnancy showed scRNAseq-based blunted immune reactivity across ventral striatal microglial subtypes^38^, but impacts of broad, physiological stress during pregnancy have not been published using scRNAseq in striatum or any brain region. Our results revealed that pathways based on striatal transcriptional changes converged across cell types, including in Drd1 and Drd2 MSNs, Sst interneurons, and microglia. With outcomes of other cell types likely limited by small cell numbers, this suggests susceptibility across cell types in the striatum to PS. Across MSN subtypes, synaptic-related pathways were upregulated. Specifically, glutamatergic synaptic transmission was repeatedly represented, suggesting upregulation of cortical glutamatergic input, in line with increased cortico-striatal connectivity in children with NDDs^5,9^. However, previous striatal glutamate studies in ASD and ADHD are inconclusive when taken together^14,15,17^, and ASD striatal molecular studies are limited.

The other most consistent theme from these scRNAseq results was downregulation of genes from translation- and ribosome-related pathways across multiple cell types. Dysregulation of translation has been implicated in NDDs, specifically local translation at axon terminals of synaptic proteins^39^. Fragile X syndrome, the most common cause of monogenic ASD and a risk for other NDDs, results from dysregulation of translation following the loss of function of the fragile X mental retardation protein (FMRP), a translational repressor^40^. Separately, mutations in ribosomal components and biogenesis factors are associated with several NDDs^41^. Evidence from human cerebral organoids shows that decreased ribosome availability during critical periods in early development impacts translation-dependent cell survival and cell fate commitment, which would have a profound, long-lasting impact on the brain^42^. Brain molecular analysis at early developmental timepoints has demonstrated similar upregulation of synapse pathways and downregulation of translation with prenatal stress^43^, and analyses in other models of NDD risk would provide more insight on the origins of striatal alterations.

Considering well-established links between PS, dorsal striatum, and ASD, we tested whether DEGs in striatal cell types after PS were overrepresented among ASD-associated genes. We found that ASD-associated genes were highly overrepresented in both Drd1 and Drd2 MSN DEGs. Previous studies have shown that ASD-associated genes are overrepresented for genes expressed in striatal neurons^18,44^, but it remains that the ASD-associated genes were more frequent among those changed by PS. This was not true of ADHD-associated genes, suggesting specificity of molecular impacts in this PS model.

Since many people with NDDs have deficits in motor learning^45^, we hypothesized that offspring in this PS model would display a deficit in rotarod-based motor learning. Contrary to that, we observed an increase in motor learning performance after PS. Notably, among mouse models of ASD, rotarod phenotypes are mixed, with both increased and decreased learning compared to controls^46^. As others have suggested, enhanced learning on rotarod could be interpreted as enhanced ability or tendency to learn a fixed motor routine, reminiscent of ASD traits.

Increased variability in time interval estimation has been reported in individuals with ASD^47^. Increased variability as well as decreased accuracy in interval timing has been shown in the valproic acid mouse model of NDDs^48^, and a shift toward earlier interval timing has been shown in a valproic acid rat model^49^. Although PS did not affect variability in interval timing here, we did observe a shift in accuracy, or mean switch time, with PS mice switching earlier. One defining characteristic of NDDs is impaired behavioral flexibility, which could contribute significantly to the shift in interval timing behavior described here. Increased impulsivity, which is also characteristic of NDDs, particularly ADHD^50,51^, could also contribute to earlier switch times.

Our multielectrode recording in mice performing the interval timing task revealed an increased proportion of MSNs that displayed time-related ramping activity during the first six seconds of interval timing switch trials by PS. We also observed an increase in absolute value PC1 score, which represents ramping activity, and absolute value MSN firing rate slope by PS, consistent with the earlier mean switch time. Since time-related ramping during this period in the trial is predictive of timing behavior,^35-37^ it follows that faster ramping would result in faster timing in the task. MSN ramping activity has been modeled to arise from cortical-striatal network characteristics^52^, and changes in cortical glutamatergic inputs onto MSNs could contribute to faster ramping MSNs. This aligns with our sequencing results, which indicated upregulated glutamatergic synaptic transmission in Drd1 and Drd2 MSNs after PS.

For statistical power and feasibility, we included only male mice in scRNAseq investigation which is a limitation. We selected males due to the higher NDD prevalence in males^53^. Although we did not observe behavioral sex differences, scRNAseq of female dorsal striatum after PS could reveal molecular level sex differences underlying shared behavioral deficits.

## Conclusions

This is the first study leveraging an scRNAseq approach to profile brain transcriptome after prenatal stress, focusing on dorsal striatum. These results revealed specific, long-term effects of *in utero* exposure to maternal stress that were previously unknown. Future work examining individual genes or pathways revealed by our sequencing data (e.g., glutamatergic synapses onto MSNs, translation, or ribosome structure in dorsal striatum) and investigating them as possible mechanistic targets for correcting functional deficits would be beneficial to translation of this study. Expansions on this work may provide potential targets for novel interventions and therapeutics for NDDs.

## Methods

### Mouse model

Wildtype CD-1 females were mated in house with GAD67-GFP^+/-^ CD-1 males (Charles River, US) to visualize striatal GABAergic neurons in later tissue analysis. Detection of vaginal plug was used to determine embryonic day (E) zero. Pregnant dams were randomly assigned to one of two groups, receiving either repetitive restraint stress or only saline injection serving as a handling control from E12 until birth. Repetitive restraint stress consisted of saline injection (intraperitoneal sterile 0.2 ml 0.9% NaCl) and then restraint in a clear plexiglass tube under bright light (13W LED bulb). Restraint stress occurred in 45-minute sessions three times daily, at approximately 08:00, 12:00, and 16:00, and control dams received only injections at these same times (n = 13 control (CTL) dams; 12 prenatal stress (PS) dams).

On postnatal day zero, litters were culled to 6-9 pups with male and female offspring as equally balanced as possible. Litters were then left undisturbed until weaning at postnatal day 21, when offspring were weaned by sex into standard housing with 2-5 littermates. Mice were maintained on a 12-hour light/dark cycle with *ad lib* access to food and water, except as noted for the operant based interval timing task described below.

All experimental procedures involving animals were performed in accordance with the University of Iowa Institutional Animal Care and Use Committee (IACUC) standards.

### Behavioral testing

GAD67-GFP^+/-^ offspring (n = 21 mice/13 litters CTL; 22 mice/12 litters PS; equal number of males and females) were tested on a series of behaviors. All tests were conducted during the light cycle. Mice were given at least 30 minutes of habituation to the room before testing for open field and rotarod. All mice started behavior at 8 or 9 weeks of age and continued testing for 8-9 weeks. Tests were conducted in the order listed below, least to most stressful.

### Open field

Open field is a standard method to assess activity level and anxiety-relevant movement. Mice were placed in a novel 1,500 cm^2^ rectangular plastic chamber with white walls and no ceiling. Mice were recorded for 30 minutes using a camera positioned above the apparatus. ANY-maze software (Stoelting Co., Wood Dale, IL) was used to track locomotor activity, with measures including total distance traveled and time spent in the center zone.

### Rotarod

Rotarod is frequently used to demonstrate motor learning, a form of procedural learning. The rotarod apparatus (Ugo Basile, Gemonio, Italy) was programmed to accelerate from 4-80 rpm over four minutes. Mice were placed on the rotating rod and latency to fall or second full spin around rod was recorded. A maximum time of five minutes was enforced. Mice were given five trials per day with at least 15 minutes between trials for three consecutive days. There were approximately 24 hours between the start of training each day.

### Interval timing switch task

We used an interval timing switch task, described in detail previously^36,54-56^. This interval timing task is a measure a rodent’s internal estimate of time, as in other interval timing tasks^54,56-60^. Briefly, mice were food restricted and trained in standard operant chambers enclosed in sound attenuating cabinets (MedAssociates, St. Albans, VT). Mice were trained to respond at either a designated short or long trial nosepoke response port after 6 or 18 seconds to receive a 20 mg sucrose reward. Trials were self-initiated and had identical cues. In a short 6-second trial, mice received a reward for the first response after 6 seconds at the designated short trial response port. In a long 18-second trial, a reward was delivered for the first response after 18 seconds at the designated response port. Since cues are identical for both trial types, the optimal strategy is to start at the short trial response port and then *switch* to the long trial response port if enough time has passed without reward. The switch response time, or the moment mice leave the short trial response port, is a measure of the rodents’ interval estimate of time. Switch times were used to calculate mean switch time (measure of interval timing accuracy) and switch time coefficient of variance (CV; measure of interval timing precision).

### Surgical procedures

A total of 8 mice (4 control and 4 prenatal stress (2 males and 2 females per group)) trained in the interval timing switch task were implanted with 16-channel microelectrode recording arrays (MicroProbes, Gaithersburg, MD) in dorsomedial striatum (DMS). Mice were anesthetized in an induction chamber using vaporized isoflurane (4%–5%), and a surgical level of anesthesia was maintained using 1.0%– 3.0% isoflurane at 120 mL/min. Under aseptic surgical conditions, the scalp was retracted, and the skull was leveled between bregma and lambda. A craniotomy was drilled above the left or right DMS and at least three craniotomies were drilled for skull screws that stabilize headcap assemblies and ground electrode arrays. Arrays consisted of 50 µm stainless steel wires in a 4×4 configuration with 250 µm between wires and rows. It was centered over the following coordinates (from bregma): AP +0.9 mm, ML ±1.4 mm, DV -2.7 mm. Cyanoacrylate (‘SloZap’ and ‘ZipKicker’, Pacer Technologies, Rancho Cucamonga, CA) and methyl methacrylate (AM Systems, Port Angeles, WA) ‘ZipKicker’ (Pacer Technologies) were used to secure arrays, skull screws, and ground wires. Mice were given at least one week to recover before reintroduction to the interval timing switch task and recording procedures.

### Neurophysiological recordings

Neuronal ensemble recordings were obtained using the multielectrode recording system (Open Ephys, Atlanta, GA). Plexon Offline Sorter (Plexon, Dallas, TX) software was used to remove artifacts and classify single neurons using principal component analysis (PCA) and waveform shape. Single neurons were defined as those 1) having a consistent waveform shape; 2) being a separable cluster in PCA space; and 3) having a consistent refractory period of at least 2 milliseconds in interspike intervals. Single units were further categorized as putative medium spiny neurons (MSNs) or striatal fast-spiking interneurons (FSIs) based on hierarchical clustering of the waveform peak-to-trough ratio and the half-peak width (*fitgmdist* and *cluster.m*) (Supplementary Fig. 1e). Only MSNs were included in subsequent analyses. Spike activity was calculated with kernel density estimates of firing rates across the interval (-4 seconds before trial start to 22 seconds after trial start) binned at 0.25 seconds, with a bandwidth of 1.

### Tissue processing and immunohistochemistry

Following dorsal striatal recordings, all mice were sacrificed to confirm histological placement of microelectrode recording arrays (Supplementary Fig. 1d). In addition, separate GAD67-GFP^+/-^ offspring were collected for neurobiological analyses between 13-17 weeks of age (n = 8-10 mice/6-8 litters CTL; 9-11 mice/8-9 litters PS; equal number of males and females). Mice were euthanized by ketamine/xylazine anesthesia before intracardiac perfusion of ice-cold phosphate buffered saline (PBS) followed by 4% paraformaldehyde (PFA). Brains were post-fixed in 4% PFA for two days before transfer to 30% sucrose in PBS. Brains were frozen in Tissue-Tek Optimal Cutting Temperature embedding medium and sectioned at 40-50 μm using a cryostat.

Unless noted below, the following procedure was used for all immunohistochemical stains: Floating sections were blocked for 1-2 hours with 10% horse serum, 0.2% Tween, and 0.1% Triton-X in PBS. Primary antibody incubation was performed overnight at room temperature (rabbit anti-GFAP;1:500, Invitrogen, Cat# PA1-10019) or 4°C (rabbit anti-Iba1; 1:500, Fujifilm Wako, Cat# 019-19741), followed by 3 washes in PBS the next day. Secondary antibody incubation (goat anti-rabbit 594; 1:500, Alexa Fluor, Cat# A-11012) was performed for 90-120 minutes, followed by an additional 3 washes in PBS. Sections were mounted in DAPI.

For Substance P, antigen retrieval was performed in 50 mM sodium citrate buffer (pH = 8) at 60°C for two hours. Primary antibody incubation was performed overnight at room temperature with rabbit anti-Substance P (1:500, Millipore, Cat# AB1566). Secondary antibody incubation was performed for 90 minutes with goat anti-rabbit 594 (1:500, Alexa Fluor, Cat# A-11012).

For Enkephalin, antigen retrieval was performed in 50 mM sodium citrate buffer (pH = 8.5) at 80°C for 30 minutes. Tissue was permeabilized for 2 hours in a PBS solution with 0.5% Triton-X and 5% horse serum. Primary antibody incubation was performed for three days at 4°C with mouse anti-Enkephalin (1:500, Santa Cruz, Cat# sc-47705), followed by 3 washes in PBS. Secondary antibody incubation was performed overnight at room temperature with goat anti-mouse 594 (1:500, Alexa Fluor, Cat# A-11005).

### Stereology

Unbiased stereology was used to quantify density of immunostained cells^61^ using a fluorescent Axiocam microscope (Zeiss, Oberkochen, Germany) coupled with StereoInvestigator software (Microbrightfield, Williston, VT). An 83 X 120 X 10 μm counting frame on a 500 X 500 μm grid was used. Every tenth serial section bilaterally was analyzed, yielding a total 7-9 sections with dorsal striatum per brain.

### Single-cell RNA sequencing

At 8-12 weeks of age, GAD67-GFP^-/-^ male offspring were euthanized by ketamine/xylazine anesthesia followed by intracardiac perfusion of artificial cerebrospinal fluid buffer (n = 4 mice/4 litters prenatal stress; n = 4 mice/3 litters control). Dorsal striatum was microdissected for single-cell suspension preparation. Buffer and single-cell suspensions were prepared as previously described^62^.

### Single cell library preparation and sequencing

Single cell libraries were generated using the Chromium Next GEM Single Cell 3’ Kit v3.1 as per manufacturer’s instructions and sequenced using the Illumina NovaSeq6000 using 100 bp paired-end reads at the University of Iowa Institute of Human Genetics Genomics Division. Cell Ranger program (10X Genomics) was used to demultiplex raw base call files and generate FASTQ files.

### Quality control, clustering and celltype identification

FASTQ files were evaluated using FastQC (v0.11.8) and MultiQC (v1.5) to determine excellent sequencing quality (Phred score > Q30). The cellranger count function from Cellranger-6.0.0 was used to align the FASTQ files to the mouse reference genome (mm10; pre-built reference mm10-2020-A obtained from the 10X genomics website) and for feature quantification. The average number of reads per cell was ∼50,000 (Supplementary Table 1). Cellranger aggr was used for data aggregation. Data was subsequently loaded into RStudio (v4.1.1, Boston, MA)^63^ and processed using Seurat (4.1.1) for normalization, data reduction, clustering, and visualization. Briefly, high quality cells defined by percentage mitochondria <10%^64^ and total feature counts >500 were retained for downstream analysis. The SCTransform function was used to normalize the data and control for confounding sources of variation such as read depth and mitochondria percentage. Based on knee plots, 15 principal components were used to create a shared nearest neighbor graph. Cells were clustered into groups by optimizing a modularity function (Louvain algorithm, resolution 0.8, 10 random starts and 10 iterations). Data was visualized using t-distributed stochastic neighbor embedding (tSNE) plots. Doublets were identified using (1.8.0)^65^ and removed from the analysis. The number of doublets was ∼2-3%, consistent with the expected number of doublets for 10X single cell RNA-Seq as per manufacturer’s guidelines. Cluster identity was determined by cell type specific marker gene expression including GABAergic neurons (*Gad1* and *Gad2*), astrocytes (*Aldoc* and *Apq4*), mature oligodendrocytes (*Mog* and *Cldn11*), oligodendrocyte precursors (*Pdgfra*), endothelia (Pecam1 and Slc2a1), microglia (*Tmem119*), pericytes (*Vtn*), endothelia (*Pecam1* and *Slc2a1)*, transient amplifying progenitors (*Pcna*, *Mki67*), neuroblasts (*Dcx*) and cholingergic neurons (*Chat*) (Supplementary Fig. 3). GABAergic neurons were further subclustered by tSNE and the identity of the subclusters were determined by cell type marker expression including *Drd1* expressing neurons, Drd2 neurons (*Drd2*, *Adora*), *Th* expressing neurons, Pthlh- and Pvalb-positive neurons (*Pthlh*^+^/*Pvalb*^+^), Pthlh-positive and Pvalb-negative (*Pthlh*^+^/*Pvalb*^-^) neurons, somatostatin-expressing neurons (*Sst*), *Cholecystokinin* expressning neurons (*Cck*), neuropeptide Y expressing (*Npy*) as previously described^66^ (Supplementary Fig. 4).

### Differential gene expression

Differential gene expression analysis was performed using NEBULA-HL^67^ as implemented in scDEpipelineR6^68^. NEBULA-HL was previously shown to provide a low false positive rate and a higher true positive rate compared to a number of other approaches^68^. Only cell types with at least 10 cells per individual mouse was used for differential gene expression (DEG) analysis. Significance testing was performed to evaluate the effect of stress condition on gene expression while controlling for batch defined by the date of collection (∼Condition + Date.of.collection). False discovery rate (FDR) was used to correct for multiple testing.

In a further analysis, ASD risk genes from the Simons Foundation Autism Research Initiative (SFARI) gene database (scores 1-3; list downloaded 01/02/2024) were tested for overrepresentation among DEGs for each cell type^69^. A curated list of candidate ADHD risk genes was also tested for overrepresentation among DEGs for each cell type. The list was compiled by integrating findings from recently published genomic studies, focusing primarily on common genetic variants associated with ADHD. Curated gene lists included those from the latest large-scale ADHD GWAS^70^, a rare CNV-based credible gene set curated by Harich *et al*.^71^, and TWAS findings from Liao *et al*.^72^ and Cabana-Dominguez *et al*.^73^. Additionally, we identified 11 ADHD GWAS studies published since 2018 through the NHGRI-EBI GWAS Catalog^74^. Genes were mapped to genome-wide significant loci using a prioritized set of mapping strategies, including Open Targets^75^ locus-to-gene (L2G) model or closest transcription start site (TSS), author-submitted annotations, and Ensembl mapping. The gene list compilation was performed using R v4.3.1, with the NHGRI-EBI GWAS Catalog accessed via REST API using the *gwasrapidd* R package^76^. To test for overrepresentation, Fisher’s Exact Test was employed in R Studio. The total population of genes was set at 20,000.

### Pathway Enrichment Analysis

For each cell type, DEGs with *p* < 0.05 after FDR correction were input into the Database for Annotation, Visualization, and Integrated Discovery (DAVID) and the Protein Analysis Through Evolutionary Relationships (PANTHER) database for functional annotation, using GOTERM_BP_DIRECT and GO-Slim Biological Process, respectively^77-79^. Resultant pathways with fewer than 10 or more than 500 genes were eliminated from the analysis.

### Statistical analyses

Data from behavioral tests and immunohistochemical measures were first analyzed using a two-way ANOVA with the variables sex and condition. Since there were no significant main effects of sex or sex by condition interactions, male and female data were combined and analyzed using two-tailed Student’s *t*-test. For repeated measures data on the rotarod, a two-way ANOVA was used with the variables of time and condition. Tests were conducted in Prism (GraphPad, San Diego, CA).

For the neurophysiological recording data, we fit a generalized linear model (GLM; *fitglme* in MATLAB) to firing rate for each neuron during interval timing switch trials described above. Neurons were classified as either “ramping” or “non-ramping”, as in prior work^36,37^, based on whether it displayed a significant, non-zero slope in firing rate from 0 to 6 seconds (*anova* in MATLAB) after correction for multiple comparisons using Benjamini-Hochberg false discovery rate. We also performed a principal component analysis of neuron activity patterns. We compared the proportion of ramping neurons, score for the first principal component, and firing rate slope between conditions using Fisher’s Exact Test (R Studio) for the former and two-tailed Student’s *t*-test (Prism, GraphPad) for the latter two.

## Data availability

At publication, the single-cell RNA sequencing data will be deposited in the Gene Expression Omnibus database.

## Acknowledgements

The authors thank the Genomics Division of the Iowa Institute of Human Genetics for their sequencing services. This work was supported by NIMH (R01 MH122485-01) to H.E.S. M.M.E. was supported by NIMH (F31 MH131259). S.V.M. and G.S.K. were supported by NIMH (T32 MH019113).

## Author contributions

Conceptualization, M.M.E., B.W.Q.H., M.A.W., N.S.N., H.E.S.; Formal Analysis, M.M.E., B.W.Q.H., M.A.W., S.V.M.; Investigation, M.M.E., B.W.Q.H., M.A.W., S.V.M., A.I.B., S.L.A., R.M.E., K.S.; Resources, G.S.K., J.J.M., N.S.N., H.E.S.; Writing - Original Draft, M.M.E., B.W.Q.H, M.A.W., A.I.B., G.S.K., H.E.S.; Writing – Review & Editing, M.M.E., M.A.W., H.E.S.; Visualization, M.M.E., B.W.Q.H., M.A.W.; Supervision, J.J.M., N.S.N., H.E.S.; Project Administration, H.E.S.; Funding acquisition, H.E.S., M.M.E., S.V.M., G.S.K.

## References

1 Han, V. X. et al. Maternal acute and chronic inflammation in pregnancy is associated with common neurodevelopmental disorders: a systematic review. Transl Psychiatry 11, 71 (2021). 10.1038/s41398-021-01198-w

2 Manzari, N., Matvienko-Sikar, K., Baldoni, F., O’Keeffe, G. W. & Khashan, A. S. Prenatal maternal stress and risk of neurodevelopmental disorders in the offspring: a systematic review and meta-analysis. Soc Psychiatry Psychiatr Epidemiol 54, 1299–1309 (2019). 10.1007/s00127-019-01745-3

3 Walder, D. J. et al. Prenatal maternal stress predicts autism traits in 6(1/2) year-old children: Project Ice Storm. Psychiatry Res 219, 353–360 (2014). 10.1016/j.psychres.2014.04.034

4 Abbott, P. W., Gumusoglu, S. B., Bittle, J., Beversdorf, D. Q. & Stevens, H. E. Prenatal stress and genetic risk: How prenatal stress interacts with genetics to alter risk for psychiatric illness. Psychoneuroendocrinology 90, 9–21 (2018). 10.1016/j.psyneuen.2018.01.019

5 Evans, M. M., Kim, J., Abel, T., Nickl-Jockschat, T. & Stevens, H. E. Developmental Disruptions of the Dorsal Striatum in Autism Spectrum Disorder. Biol Psychiatry 95, 102–111 (2024). 10.1016/j.biopsych.2023.08.015

6 Hong, Y. N., Hwang, H., Hong, J. & Han, D. H. Correlations between developmental trajectories of brain functional connectivity, neurocognitive functions, and clinical symptoms in patients with attention-deficit hyperactivity disorder. J Psychiatr Res 173, 347-354 (2024). 10.1016/j.jpsychires.2024.03.021

7 Hoogman, M. et al. Subcortical brain volume differences in participants with attention deficit hyperactivity disorder in children and adults: a cross-sectional mega-analysis. Lancet Psychiatry 4, 310–319 (2017). 10.1016/S2215-0366(17)30049-4

8 Langen, M., Durston, S., Staal, W. G., Palmen, S. J. & van Engeland, H. Caudate nucleus is enlarged in high-functioning medication-naive subjects with autism. Biol Psychiatry 62, 262–266 (2007). 10.1016/j.biopsych.2006.09.040

9 Di Martino, A., et al. Aberrant striatal functional connectivity in children with autism. Biol Psychiatry 69, 847-856 (2011). 10.1016/j.biopsych.2010.10.029

10 Brandenburg, C. et al. Increased Dopamine Type 2 Gene Expression in the Dorsal Striatum in Individuals With Autism Spectrum Disorder Suggests Alterations in Indirect Pathway Signaling and Circuitry. Front Cell Neurosci 14, 577858 (2020). 10.3389/fncel.2020.577858

11 Portmann, T. et al. Behavioral abnormalities and circuit defects in the basal ganglia of a mouse model of 16p11.2 deletion syndrome. Cell Rep 7, 1077–1092 (2014). 10.1016/j.celrep.2014.03.036

12 Kuo, H. Y. & Liu, F. C. Pathological alterations in striatal compartments in the human brain of autism spectrum disorder. Mol Brain 13, 83 (2020). 10.1186/s13041-020-00624-2

13 Kuo, H. Y. & Liu, F. C. Valproic acid induces aberrant development of striatal compartments and corticostriatal pathways in a mouse model of autism spectrum disorder. FASEB J 31, 4458–4471 (2017). 10.1096/j.201700054R

14 Carey, C. et al. From bench to bedside: The mGluR5 system in people with and without Autism Spectrum Disorder and animal model systems. Transl Psychiatry 12, 395 (2022). 10.1038/s41398-022-02143-1

15 Hollestein, V. et al. Developmental changes in fronto-striatal glutamate and their association with functioning during inhibitory control in autism spectrum disorder and obsessive compulsive disorder. Neuroimage Clin 30, 102622 (2021). 10.1016/j.nicl.2021.102622

16 Oron, O. et al. Gene network analysis reveals a role for striatal glutamatergic receptors in dysregulated risk-assessment behavior of autism mouse models. Transl Psychiatry 9, 257 (2019). 10.1038/s41398-019-0584-5

17 Sudre, G. et al. Mapping the cortico-striatal transcriptome in attention deficit hyperactivity disorder. Mol Psychiatry 28, 792–800 (2023). 10.1038/s41380-022-01844-9

18 Fuccillo, M. V. Striatal Circuits as a Common Node for Autism Pathophysiology. Front Neurosci 10, 27 (2016). 10.3389/fnins.2016.00027

19 Schwabe, L., Bohbot, V. D. & Wolf, O. T. Prenatal stress changes learning strategies in adulthood. Hippocampus 22, 2136–2143 (2012). 10.1002/hipo.22034

20 Diaz, R., Ogren, S. O., Blum, M. & Fuxe, K. Prenatal corticosterone increases spontaneous and d-amphetamine induced locomotor activity and brain dopamine metabolism in prepubertal male and female rats. Neuroscience 66, 467–473 (1995). 10.1016/0306-4522(94)00605-5

21 Arzuaga, A. L., Edmison, D. D., Mroczek, J., Larson, J. & Ragozzino, M. E. Prenatal stress and fluoxetine exposure in mice differentially affect repetitive behaviors and synaptic plasticity in adult male and female offspring. Behav Brain Res 436, 114114 (2023). 10.1016/j.bbr.2022.114114

22 Schroeder, R. et al. Maternal P7C3-A20 Treatment Protects Offspring from Neuropsychiatric Sequelae of Prenatal Stress. AnCoxid Redox Signal 35, 511–530 (2021). 10.1089/ars.2020.8227

23 Converse, A. K. et al. Prenatal stress induces increased striatal dopamine transporter binding in adult nonhuman primates. Biol Psychiatry 74, 502–510 (2013). 10.1016/j.biopsych.2013.04.023

24 Berger, M. A., Barros, V. G., Sarchi, M. I., Tarazi, F. I. & Antonelli, M. C. Long-term effects of prenatal stress on dopamine and glutamate receptors in adult rat brain. Neurochem Res 27, 1525–1533 (2002). 10.1023/a:1021656607278

25 Saunders, A. et al. Molecular Diversity and Specializations among the Cells of the Adult Mouse Brain. Cell 174, 1015–1030 e1016 (2018). 10.1016/j.cell.2018.07.028

26 Martin, A., et al. A Spatiomolecular Map of the Striatum. Cell Rep 29, 4320-4333 e4325 (2019). 10.1016/j.celrep.2019.11.096

27 Anderson, A. G., Kulkarni, A., Harper, M. & Konopka, G. Single-Cell Analysis of Foxp1-Driven Mechanisms Essential for Striatal Development. Cell Rep 30, 3051–3066 e3057 (2020). 10.1016/j.celrep.2020.02.030

28 Gokce, O. et al. Cellular Taxonomy of the Mouse Striatum as Revealed by Single-Cell RNA-Seq. Cell Rep 16, 1126–1137 (2016). 10.1016/j.celrep.2016.06.059

29 Lee, H. et al. Cell Type-Specific Transcriptomics Reveals that Mutant Huntingtin Leads to Mitochondrial RNA Release and Neuronal Innate Immune Activation. Neuron 107, 891–908 e898 (2020). 10.1016/j.neuron.2020.06.021

30 Kim, J. et al. A chromosome region linked to neurodevelopmental disorders acts in distinct neuronal circuits in males and females to control locomotor behavior. bioRxiv (2024). 10.1101/2024.05.17.594746

31 Kim, H. et al. Transcriptomic analysis in the striatum reveals the involvement of Nurr1 in the social behavior of prenatally valproic acid-exposed male mice. Transl Psychiatry 12, 324 (2022). 10.1038/s41398-022-02056-z

32 Yoo, T., Yoo, Y. E., Kang, H. & Kim, E. Age, brain region, and gene dosage-differential transcriptomic changes in Shank3-mutant mice. Front Mol Neurosci 15, 1017512 (2022). 10.3389/fnmol.2022.1017512

33 Yoo, Y. E., Yoo, T., Kang, H. & Kim, E. Brain region and gene dosage-differential transcriptomic changes in Shank2-mutant mice. Front Mol Neurosci 15, 977305 (2022). 10.3389/fnmol.2022.977305

34 Wichterle, H., Turnbull, D. H., Nery, S., Fishell, G. & Alvarez-Buylla, A. In utero fate mapping reveals distinct migratory pathways and fates of neurons born in the mammalian basal forebrain. Development 128, 3759–3771 (2001). 10.1242/dev.128.19.3759

35 Emmons, E. B. et al. Rodent Medial Frontal Control of Temporal Processing in the Dorsomedial Striatum. J Neurosci 37, 8718–8733 (2017). 10.1523/JNEUROSCI.1376-17.2017

36 Bruce, R. A. et al. Experience-related enhancements in striatal temporal encoding. Eur J Neurosci 54, 5063–5074 (2021). 10.1111/ejn.15344

37 Bruce, R. A. et al. Complementary cognitive roles for D2-MSNs and D1-MSNs in interval timing. bioRxiv (2024). 10.1101/2023.07.25.550569

38 Hayes, L. N. et al. Prenatal immune stress blunts microglia reactivity, impairing neurocircuitry. Nature 610, 327–334 (2022). 10.1038/s41586-022-05274-z

39 Joo, Y. & Benavides, D. R. Local Protein Translation and RNA Processing of Synaptic Proteins in Autism Spectrum Disorder. Int J Mol Sci 22 (2021). 10.3390/ijms22062811

40 Scarpin, M. R., Pastore, B., Tang, W. & Kearse, M. G. Characterization of ribosome stalling and no-go mRNA decay stimulated by the fragile X protein, FMRP. J Biol Chem 300, 107540 (2024). 10.1016/j.jbc.2024.107540

41 Hetman, M. & Slomnicki, L. P. Ribosomal biogenesis as an emerging target of neurodevelopmental pathologies. J Neurochem 148, 325–347 (2019). 10.1111/jnc.14576

42 Ni, C. et al. An inappropriate decline in ribosome levels drives a diverse set of neurodevelopmental disorders. bioRxiv (2024). 10.1101/2024.01.09.574708

43 Verosky, D. C., H. J.; Gur, T. L. in *Neuroscience MeeCng Planner*. Online (Society for Neuroscience).

44 Chang, J., Gilman, S. R., Chiang, A. H., Sanders, S. J. & Vitkup, D. Genotype to phenotype relationships in autism spectrum disorders. Nat Neurosci 18, 191–198 (2015). 10.1038/nn.3907

45 Dowell, L. R., Mahone, E. M. & Mostofsky, S. H. Associations of postural knowledge and basic motor skill with dyspraxia in autism: implication for abnormalities in distributed connectivity and motor learning. Neuropsychology 23, 563–570 (2009). 10.1037/a0015640

46 Cording, K. R. & Bateup, H. S. Altered motor learning and coordination in mouse models of autism spectrum disorder. Front Cell Neurosci 17, 1270489 (2023). 10.3389/fncel.2023.1270489

47 Isaksson, S. et al. Is there a generalized timing impairment in Autism Spectrum Disorders across time scales and paradigms? J Psychiatr Res 99, 111–121 (2018). 10.1016/j.jpsychires.2018.01.017

48 Acosta, J. et al. Deficits in temporal processing in mice prenatally exposed to Valproic Acid. Eur J Neurosci 47, 619–630 (2018). 10.1111/ejn.13621

49 DeCoteau, W. E. & Fox, A. E. Timing and Intertemporal Choice Behavior in the Valproic Acid Rat Model of Autism Spectrum Disorder. J AuCsm Dev Disord 52, 2414–2429 (2022). 10.1007/s10803-021-05129-y

50 Lyvers, M., Dark, S., Jaguru, I. & Thorberg, F. A. Adult symptoms of ASD and ADHD in relation to alcohol use: Potential roles of transdiagnostic features. Alcohol 120, 109–117 (2024). 10.1016/j.alcohol.2024.03.011

51 Grant, J. E. & Chamberlain, S. R. Autistic traits in young adults who gamble. CNS Spectr 21, 1–6 (2020). 10.1017/S1092852920001571

52 Ponzi, A. & Wickens, J. Ramping activity in the striatum. Front Comput Neurosci 16, 902741 (2022). 10.3389/fncom.2022.902741

53 Maenner, M. J. et al. Prevalence and Characteristics of Autism Spectrum Disorder Among Children Aged 8 Years - Autism and Developmental Disabilities Monitoring Network, 11 Sites, United States, 2020. MMWR Surveill Summ 72, 1-14 (2023). 10.15585/mmwr.ss7202a1

54 Larson, T. et al. Mice expressing P301S mutant human tau have deficits in interval timing. Behav Brain Res 432, 113967 (2022). 10.1016/j.bbr.2022.113967

55 Weber, M. A. et al. Alpha-synuclein pre-formed fibrils injected into prefrontal cortex primarily spread to cortical and subcortical structures and lead to isolated behavioral symptoms. bioRxiv (2023). 10.1101/2023.01.31.526365

56 Stutt, H. R. et al. Sex similarities and dopaminergic differences in interval timing. Behav Neurosci 138, 85–93 (2024). 10.1037/bne0000577

57 Balci, F. et al. Interval timing in genetically modified mice: a simple paradigm. Genes Brain Behav 7, 373–384 (2008). 10.1111/j.1601-183X.2007.00348.x

58 Tosun, T., Gur, E. & Balci, F. Mice plan decision strategies based on previously learned time intervals, locations, and probabilities. *Proc Natl Acad Sci U S A* **113**, 787-792 (2016). 10.1073/pnas.1518316113

59 Narayanan, N. S., Land, B. B., Solder, J. E., Deisseroth, K. & DiLeone, R. J. Prefrontal D1 dopamine signaling is required for temporal control. Proc Natl Acad Sci U S A 109, 20726–20731 (2012). 10.1073/pnas.1211258109

60 Emmons, E. et al. Temporal Learning Among Prefrontal and Striatal Ensembles. Cereb Cortex Commun 1, tgaa058 (2020). 10.1093/texcom/tgaa058

61 Peterson, D. A. Quantitative histology using confocal microscopy: implementation of unbiased stereology procedures. Methods 18, 493–507 (1999). 10.1006/meth.1999.0818

62 Hing, B. et al. Single Cell Transcriptome of Stress Vulnerability Network in mouse Prefrontal Cortex. bioRxiv (2023). 10.1101/2023.05.14.540705

63 RStudio: Integrated Development for R (RStudio, PBC, Boston, MA, 2020).

64 Osorio, D. & Cai, J. J. Systematic determination of the mitochondrial proportion in human and mice tissues for single-cell RNA-sequencing data quality control. BioinformaCcs 37, 963–967 (2021). 10.1093/bioinformatics/btaa751

65 Germain, P. L., Lun, A., Garcia Meixide, C., Macnair, W. & Robinson, M. D. Doublet identification in single-cell sequencing data using scDblFinder. F1000Res 10, 979 (2021). 10.12688/f1000research.73600.2

66 Munoz-Manchado, A. B. et al. Diversity of Interneurons in the Dorsal Striatum Revealed by Single-Cell RNA Sequencing and PatchSeq. Cell Rep 24, 2179–2190 e2177 (2018). 10.1016/j.celrep.2018.07.053

67 He, L. et al. NEBULA is a fast negative binomial mixed model for differential or co-expression analysis of large-scale multi-subject single-cell data. Commun Biol 4, 629 (2021). 10.1038/s42003-021-02146-6

68 Gagnon, J. et al. Recommendations of scRNA-seq Differential Gene Expression Analysis Based on Comprehensive Benchmarking. Life (Basel) 12 (2022). 10.3390/life12060850

69 Banerjee-Basu, S. & Packer, A. SFARI Gene: an evolving database for the autism research community. Dis Model Mech 3, 133–135 (2010). 10.1242/dmm.005439

70 Demontis, D., et al. Genome-wide analyses of ADHD identify 27 risk loci, refine the genetic architecture and implicate several cognitive domains. Nat Genet 55, 198-208 (2023). 10.1038/s41588-022-01285-8

71 Harich, B. et al. From Rare Copy Number Variants to Biological Processes in ADHD. Am J Psychiatry 177, 855–866 (2020). 10.1176/appi.ajp.2020.19090923

72 Liao, C. et al. Transcriptome-wide association study of attention deficit hyperactivity disorder identifies associated genes and phenotypes. Nat Commun 10, 4450 (2019). 10.1038/s41467-019-12450-9

73 Cabana-Dominguez, J. et al. Transcriptomic risk scores for attention deficit/hyperactivity disorder. Mol Psychiatry 28, 3493–3502 (2023). 10.1038/s41380-023-02200-1

74 Sollis, E. et al. The NHGRI-EBI GWAS Catalog: knowledgebase and deposition resource. Nucleic Acids Res 51, D977–D985 (2023). 10.1093/nar/gkac1010

75 Mountjoy, E. et al. An open approach to systematically prioritize causal variants and genes at all published human GWAS trait-associated loci. Nat Genet 53, 1527–1533 (2021). 10.1038/s41588-021-00945-5

76 Magno, R. & Maia, A. T. gwasrapidd: an R package to query, download and wrangle GWAS catalog data. BioinformaCcs 36, 649–650 (2020). 10.1093/bioinformatics/btz605

77 Mi, H. et al. PANTHER version 11: expanded annotation data from Gene Ontology and Reactome pathways, and data analysis tool enhancements. Nucleic Acids Res 45, D183–D189 (2017). 10.1093/nar/gkw1138

78 Huang da, W., Sherman, B. T. & Lempicki, R. A. Systematic and integrative analysis of large gene lists using DAVID bioinformatics resources. Nat Protoc 4, 44–57 (2009). 10.1038/nprot.2008.211

79 Sherman, B. T., et al. DAVID: a web server for functional enrichment analysis and functional annotation of gene lists (2021 update). Nucleic Acids Res 50, W216-W221 (2022). 10.1093/nar/gkac194

