## Supplementary Figures 1-5 for "Long-term, cell type-specific effects of prenatal stress on dorsal striatum and relevant behaviors in mice"

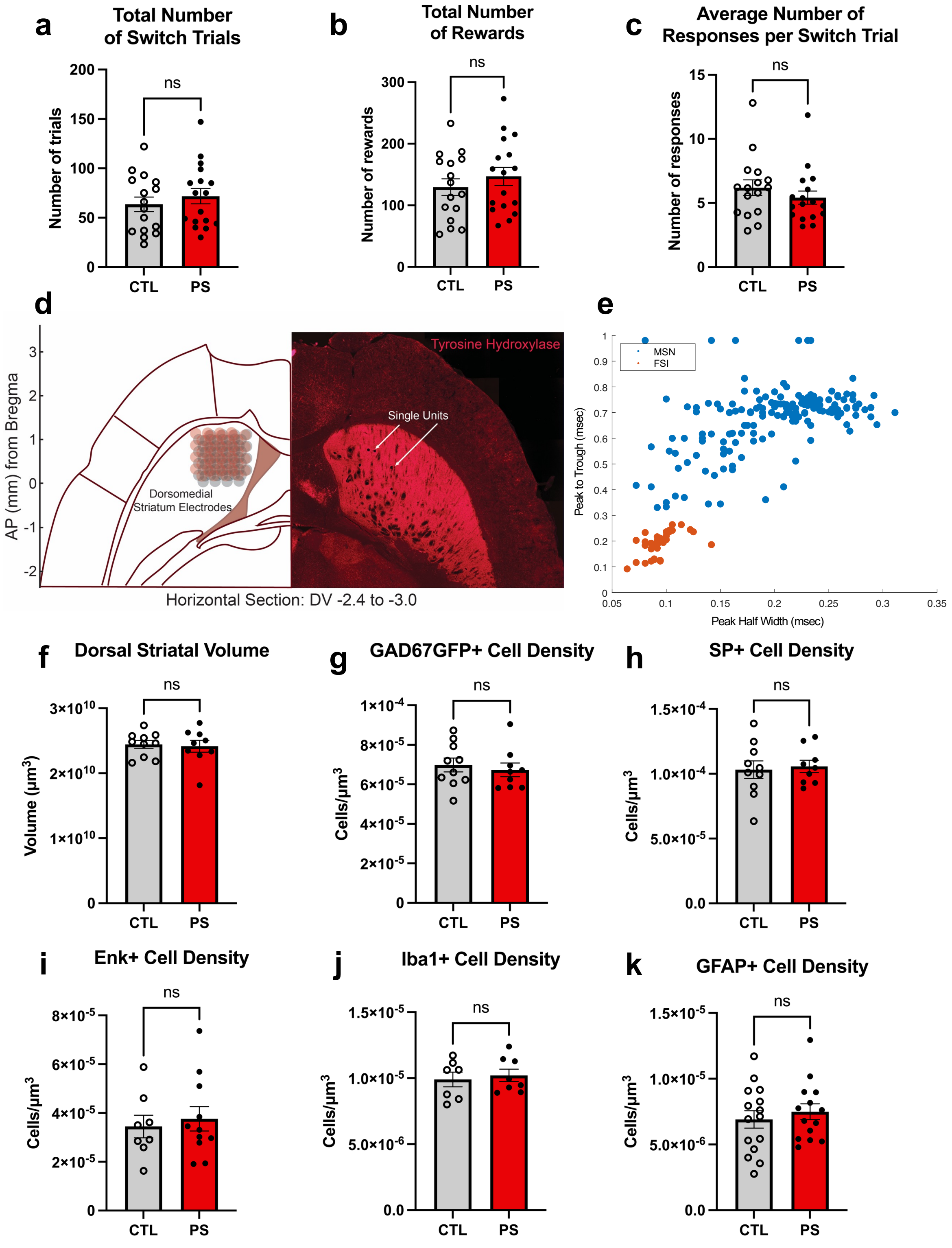


**Supplementary Figure 1: Additional behavioral measures and immunofluorescent analysis of cell types in dorsal striatum.** Prenatal stress (PS) did not affect **(a)** the total number of switch trials completed over the testing days, **(b)** the total number of rewards received over the testing days, or **(c)** the average number of nosepoke responses per switch trial in the interval timing switch task. (**d**) Immunohistochemistry for tyrosine hydroxylase in horizontal sections of the brain confirmed correct placement of the electrodes in dorsomedial striatum. (**e**) Categorization of putative medium spiny neurons (MSNs) or striatal fast-spiking interneurons (FSIs) based on hierarchical clustering of the waveform peak-to-trough ratio and the half-peak width. There were no PS effects on **(f)** dorsal striatal volume, **(g)** GAD67GFP+ (glutamic acid decarboxylase green fluorescent protein) cell density in the dorsal striatum, a marker of GABAergic neurons, **(h)** SP+ (substance P) cell density in dorsal striatum, a marker of Drd1 MSNs, **(i)** Enk+ (enkephalin) cell density in dorsal striatum, a marker of Drd2 MSNs, (**j**) Iba1+ (ionized calcium-binding adaptor molecule 1) cell density in dorsal striatum, a marker of microglia, or (**k**) GFAP+ (glial fibrillary acidic protein) cell density in dorsal striatum, a marker of astrocytes. ns = non-significant.

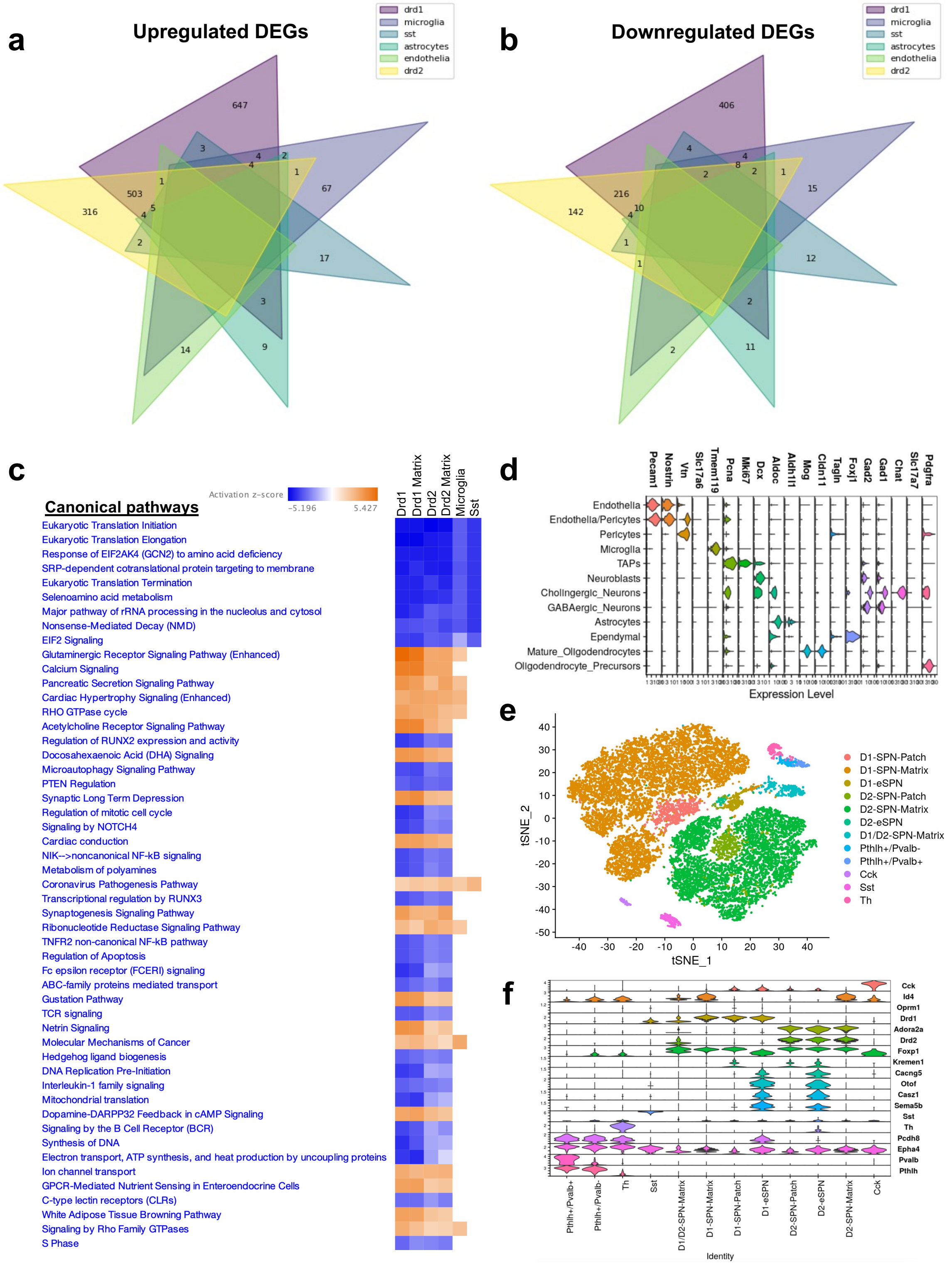


**Supplementary Figure 2: Differential gene expression, pathway analysis results, and cell clustering from scRNAseq. (a)** Several cell types in dorsal striatum had overlapping upregulated differentially expressed genes (DEGs) after prenatal stress (PS). **(b)** Several cell types in dorsal striatum had overlapping downregulated DEGs after PS. **(c)** Canonical pathway analysis of DEGs after PS were relatively similar across subpopulations of medium spiny neurons, microglia, and somatostatin (Sst) interneurons. **(d)** Expression levels of cell type marker genes for each striatal cell subtype. **(e)** T-distributed stochastic neighbor embedding (tSNE) visualization of only GABAergic neurons from original analysis (n = 19,530) from 8 samples (n = 4 control, 4 prenatal stress). **(f)** Expression levels of cell type marker genes for each GABAergic neuron subtype.

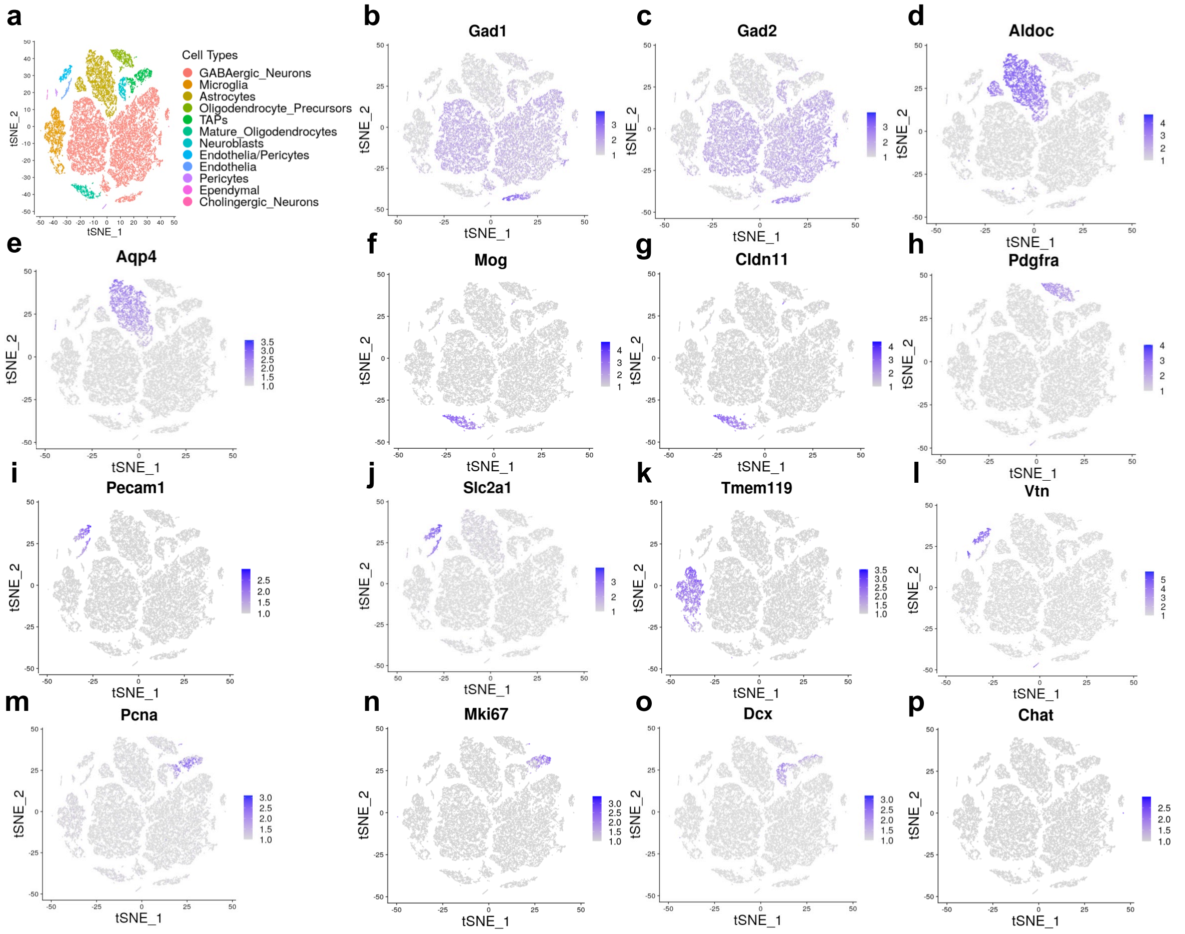


**Supplementary Figure 3: Cell type identification of clusters.** Data were clustered by T-distributed stochastic neighbor embedding (tSNE) and the identity of each cluster was determined using cell type specific markers. **(a)** Cell type identity of clusters. Cell type markers for GABAergic neurons; **(b)** *Gad1* and **(c)** *Gad2*, astrocytes; **(d)** *Aldoc* and **(e)** *Apq4*, mature oligodendrocytes; **(f)** *Mog* and **(g)** Cldn11, oligodendrocyte precursor; **(h)** *Pdgfra*, endothelia; **(i)** *Pecam1* and **(j)** *Slc2a1*, microglia; **(k)** *Tmem119*, pericytes; **(l)** *Vtn*, transient amplifying progenitors; **(m)** *Pcna* and **(n)** *Mki67*, neuroblasts; **(o)** *Dcx,* and cholingergic neurons **(p)** *Chat*.

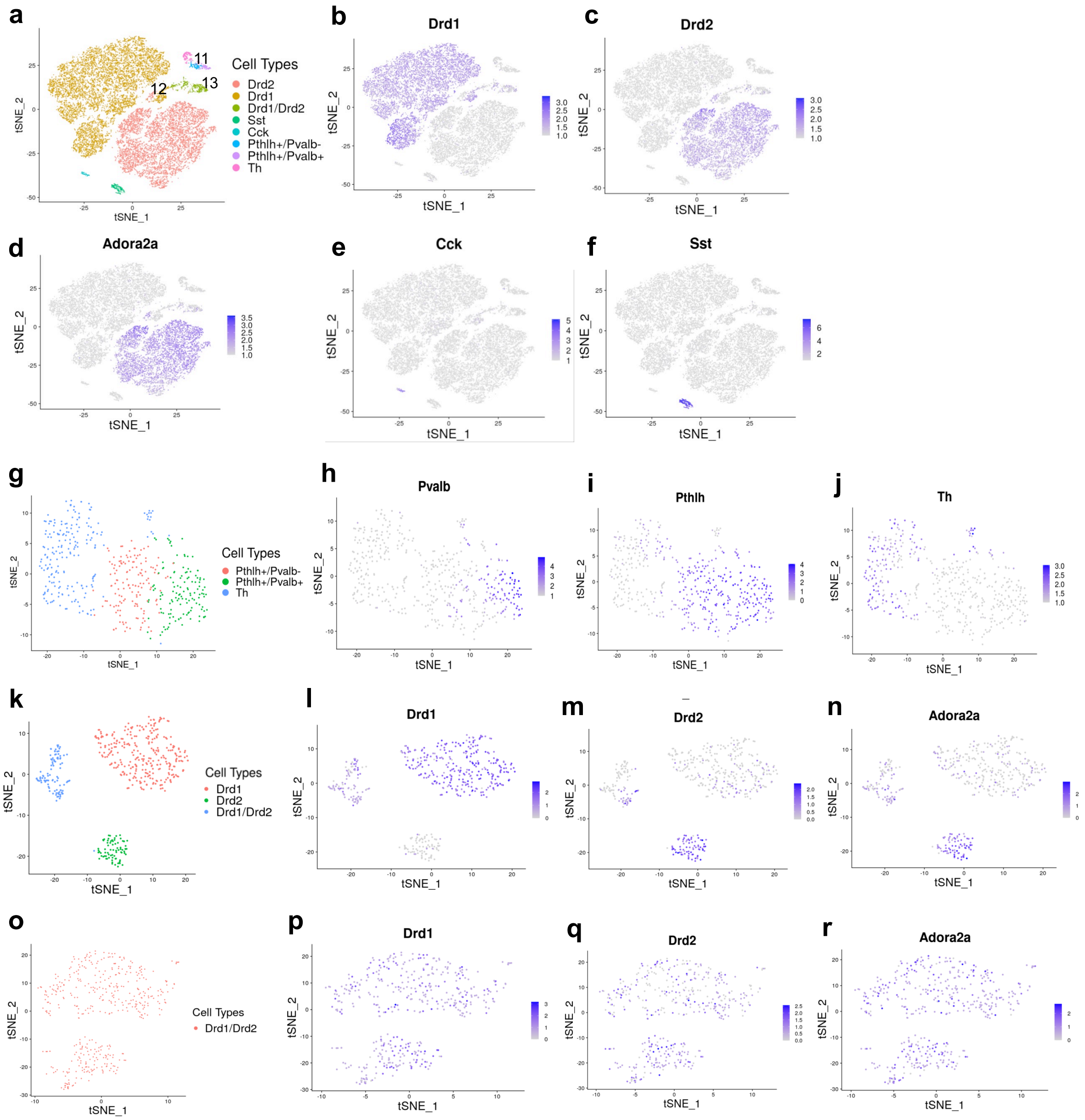


**Supplementary Figure 4: Cell type identification of sub-clustering of GABAergic neurons.** GABAergic neurons were sub-clustered by T-distributed stochastic neighbor embedding (tSNE) and the identity of each cluster was determined using cell type specific markers. **(a)** Cell type identity of clusters. Cell type markers for Drd1 neurons; **(b)** *Drd1*, Drd2 neurons; **(c)** *Drd2* and **(d)** *Adora2a*, Cholecystokinin expressing neurons; **(e)** *Cck* and Somatostatin expression neurons;

**(f)** *Sst*. Cluster 11 was further sub-clustered and the identity of each subcluster was determined using cell type specific markers. **(g)** Cell type identity of clusters. Cell type marker expression for **(h)** *Pvalb*, **(i)** *Pthlh* and **(j)** *Th*. Cluster 12 was further sub-clustered and the identity of each subcluster was determined using cell type specific markers. **(k)** Cell type identity of clusters. Cell type marker expression for Drd1; **(l)** *Drd1*, and Drd2; **(m)** *Drd2* and **(n)** *Adora2a*. Cluster 13 was further sub-clustered and the identity of each subcluster was determined using cell type specific markers. **(o)** Cell type identity of clusters. Cell type marker expression for Drd1; **(p)** *Drd1*, and Drd2; **(q)** *Drd2* and **(r)** *Adora2a*.

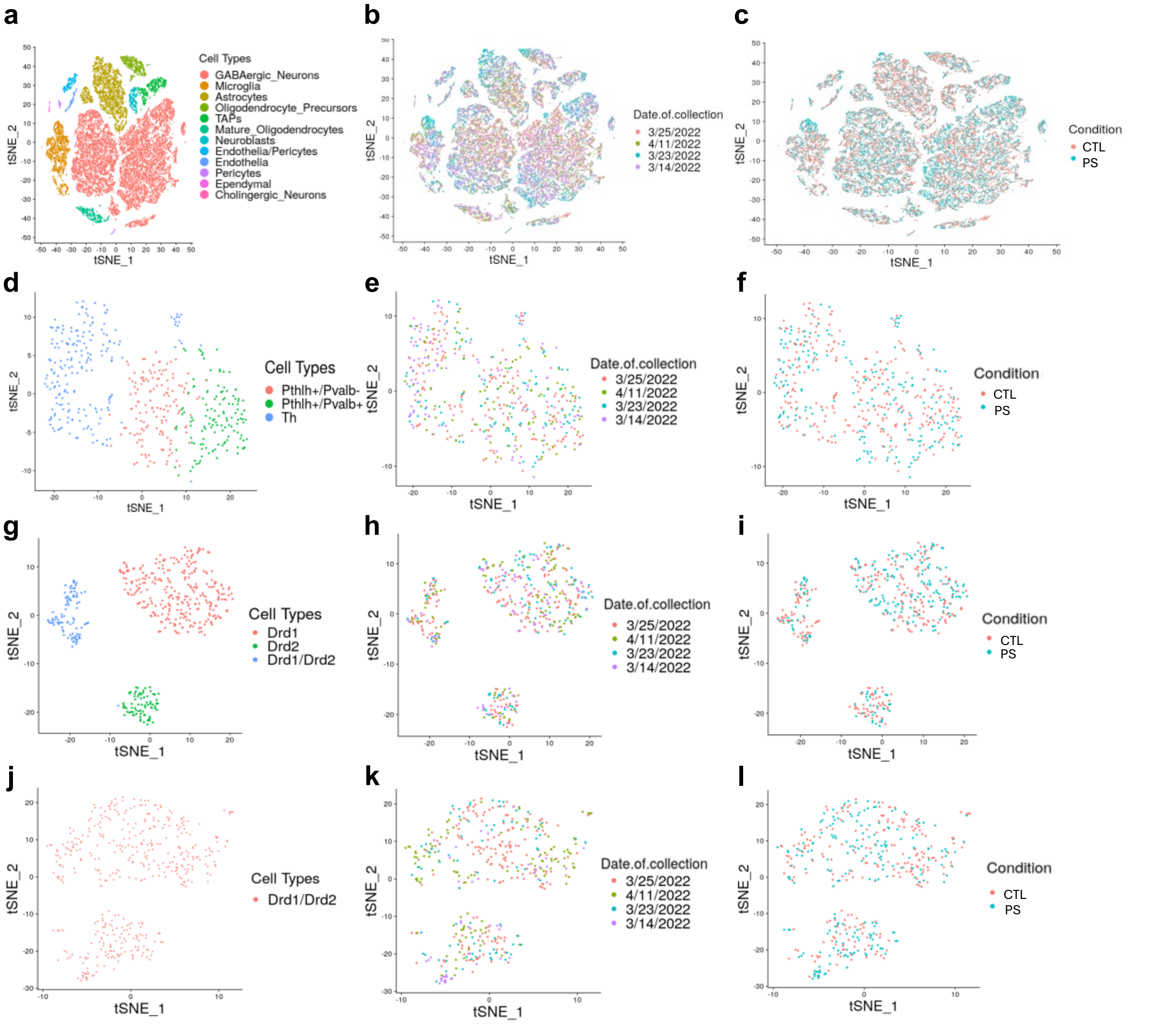


**Supplementary Figure 5: Effect of variables on tSNE clustering.** General cell type identity of clusters for **(a)** general cell types, **(d)** subcluster 11, **(g)** subcluster 12 and **(j)** subcluster 13. **(b, e, h, and k)** Overlap of batch (i.e. date of collection) and **(c, f, i, and l)** treatment condition on the respective clusters. CTL; control, PS; prenatal stress.
